# *Aspergillus fumigatus ffmA* encodes a C_2_H_2_-containing transcriptional regulator that modulates azole resistance and is required for normal growth

**DOI:** 10.1101/2021.11.17.469059

**Authors:** Sanjoy Paul, Paul Bowyer, Michael Bromley, W. Scott Moye-Rowley

## Abstract

The production of a collection of disruption mutant strains corresponding to a large number of transcription factors from the filamentous fungal pathogen *Aspergillus fumigatus* has permitted rapid identification of transcriptional regulators involved in a range of different processes. Here we characterize a gene designated *ffmA* (favors fermentative metabolism) as an C_2_H_2_-containing transcription factor that is required for azole drug resistance and normal growth. Loss of *ffmA* caused cells to exhibit significant defects in growth, either under untreated or azole-challenged conditions. Loss of FfmA caused a reduction in expression of the AbcG1 ATP-binding cassette transporter, previousy shown to contribute to azole resistance. Strikingly, overproduction of the AtrR transcription factor gene restored a wild-type growth phenotype to a *ffmAΔ* strain. Overexpression of AtrR also suppressed the defect in AbcG1 expression caused by loss of FfmA. Replacement of the *ffmA* promoter with a doxycycline-repressible promoter restored near normal growth in the absence of doxycycline. Finally, chromatin immunoprecipitation experiments indicated that FfmA bound to its own promoter as well as to the *abcG1* promoter. These data imply that FfmA and AtrR interact both with respect to *abcG1* expression and also more broadly to regulate hyphal growth.

**Importance:** Infections associated with azole-resistant forms of the primary human pathogen, *Aspergillus fumigatus*, are associated with poor outcomes in patient populations. This makes analysis of the mechanisms underlying azole resistance of *A. fumigatus* a high priority. In this work, we describe characterization of a gene designated *ffmA* that encodes a sequence-specific transcriptional regulator. We identified *ffmA* in a screen of a collection of gene disruption mutant strains made in *A. fumigatus*. Loss of *ffmA* caused sensitivity to azole drugs and also a large reduction in normal growth. We found that overproduction of the AtrR transcription factor was able to restore growth to *ffmA* null cells. We provide evidence that FfmA can recognize promoters of genes involved in azole resistance as well as the *ffmA* promoter itself. Our data indicate that FfmA and AtrR interact to support azole resistance and normal growth.

## Introduction

*Aspergillus fumigatus* is the primary filamentous fungal pathogen in humans (reviewed in (1)). Infections caused by *A. fumigatus* usually respond to azole antifungal drugs but isolates that show dramatic decreases in susceptibility are appearing with increased frequency around the world (2). Azole drugs, like voriconazole, target the enzymatic product of the *cyp51A* gene called lanosterol α-14 demethylase (3). One of the best-known alleles associated with azole resistance in *A. fumigatus* consists of a compound mutation corresponding to a duplication of a 34 bp region in the *cyp51A* promoter along with a single amino acid substitution in the encoded enzyme (4). This allele is referred to as TR34 (34 bp promoter duplication) L98H *cyp51A*. Together, these two alterations cause a dramatic increase in voriconazole MIC. Infections associated with this *cyp51A* allele are associated with a significantly worse outcome in particular populations (5) and make understanding regulation of *cyp51A* in particular and azole resistance in general of high priority in this pathogen.

While mutations associated with the *cyp51A* gene are found with high frequency in azole-resistant *A. fumigatus* isolates, more recent studies have demonstrated that not all azole resistance in this organism can be explained by mutations at *cyp51A* (reviewed in (6, 7)). Mutations in the gene encoding HMG CoA reductase have been described that lead to strong decreases in voriconazole susceptibility with a wild-type *cyp51A* locus present (8, 9). Additionally, an alteration in the *hapE* gene encoding a subunit of the CCAAT-binding complex also produced a strain with an elevated voriconazole MIC (10). Loss of normal *hapE* function leads to a large increase in *cyp51A* expression with associated elevation of azole resistance (11). Coupled with the well-described increase in *cyp51A* expression caused by the presence of the TR34 promoter (11, 12), these data illustrate the critical contribution of transcriptional regulation to azole resistance.

An additional *non-cyp51A* mode of azole resistance is illustrated by the overproduction of the ATP-binding cassette (ABC) transporter AbcG1 (aka Cdr1B/abcC). Clinical isolates have been described in which elevated transcription of the gene encoding this membrane protein leads to enhanced azole resistance (13). The presence of the *abcG1* gene is required for the normal increase in voriconazole resistance seen in strains containing the TR34 L98H form of *cyp51A* demonstrating the role of this ABC transporter even in the presence of a strongly resistant allele of *cyp51A* (14). We and others have found a transcription factor designated AtrR (ABC transporter regulator) that is required for high level transcription of both *abcG1* and *cyp51A* (15, 16). Transcriptional control of *cyp51A* is a key contributor to azole resistance but much remains to be uncovered concerning the range of transcriptional inputs to drug resistance in *A. fumigatus*.

We have previously described a collection of disruption mutants in genes that encode transcription factors (17). The first large-scale screening of this mutant collection employed itraconazole and led to the identification of the negative transcriptional regulator *nctA/B*. This mutant library also confirmed that disruption mutants in either the sterol gene regulator *srbA* (18) or *atrR* caused large increases in itraconazole susceptibility. We counterscreened these itraconazole susceptible mutant strains for effects on expression of the AbcG1 ABC transporter using an antibody directed against this membrane protein (14). A poorly characterized gene called *rfeC* (recently renamed *ffmA*) (19) was identified that caused a large decrease in AbcG1 expression. The *ffmA* gene encodes a protein containing a C_2_H_2_-containing sequence-specific DNA-binding domain (20). We also found that *ffmA* null mutants have a large defect in normal growth along with the predicted azole hypersensitivity. Interestingly, overexpression of the AtrR transcription factor was able to suppress both the growth defect of an *ffmA* null mutant and restore AbcG1 expression. These data indicate the presence of a genetic interaction between AtrR and FfmA.

## Results

### Screening the transcription factor disruption mutant library for genes that affect azole resistance

Using a previously-described transcription factor disruption mutant library (17), we identified additional mutant strains that exhibited increased susceptibility to itraconazole. These individual strains were grown from each library isolate, whole cell protein extracts prepared and these analyzed by western blotting using an anti-AbcG1 antiserum (21). The results of this western blot comparison are shown in Figure 1.

**Figure 1.**
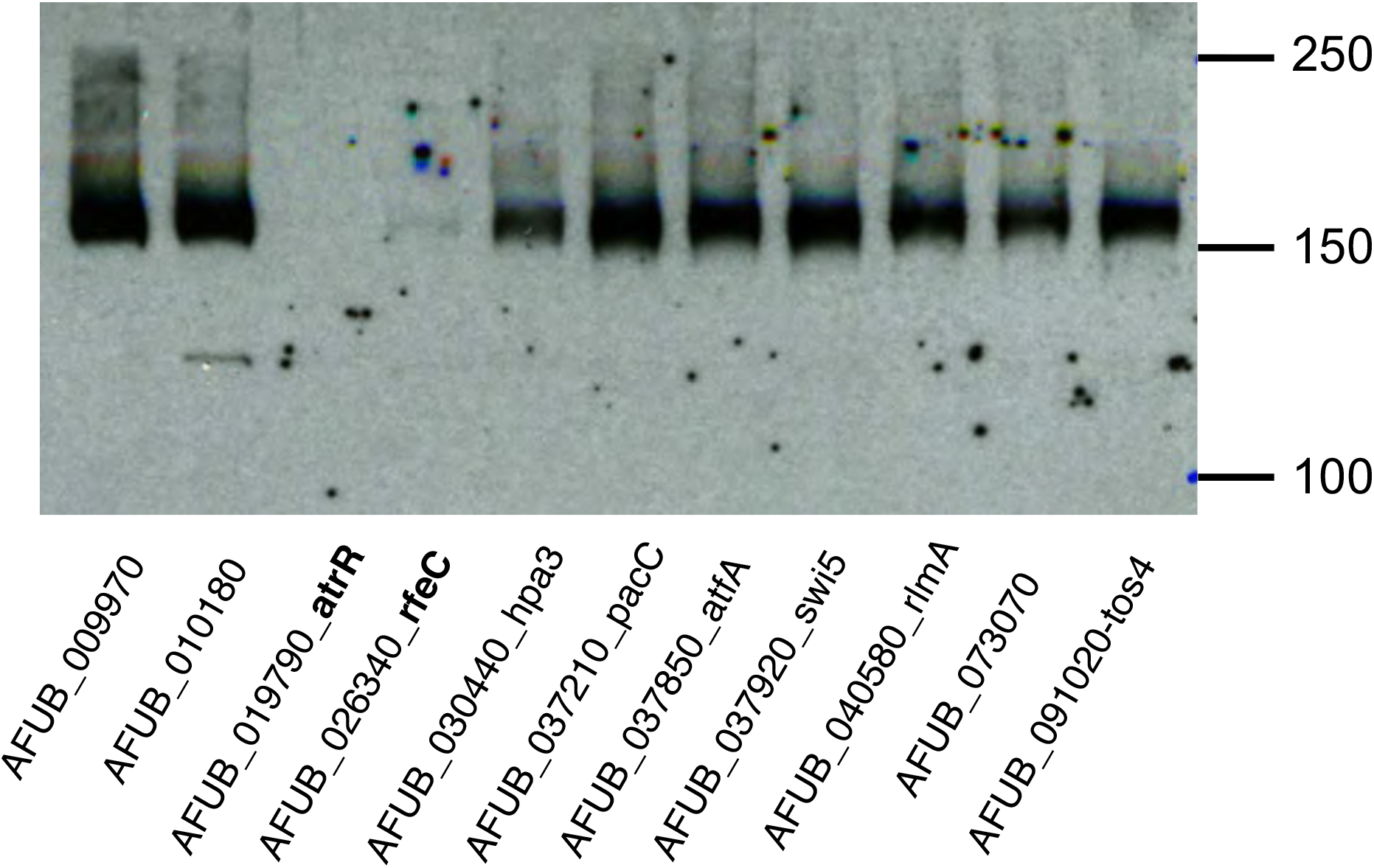
Western blot of AbcG1 levels in itraconazole susceptible transcription factor disruption strains. The indicated strains were grown to mid-log phase and whole cell protein extracts prepared. These extracts were resolved on SDS-PAGE and then analyzed by western blotting using an anti-AbcG1 polyclonal antiserum (21). To ensure equivalent loading of protein extracts, the membrane was stained with Ponceau S dye after transfer.

We found clear and reproducible reduction in the level of AbcG1 in strains lacking either the *atrR* or AFUB_026340 (*Afu2g10550*) genes. This latter gene has recently been characterized as a locus enhancing expression of genes involved fermentation (favors fermentative metabolism: *ffmA* (19)) and we will use this designation here. The *ffmA* gene has also been characterized as *rfeC* (regulator of *FLO11*) on the basis of activating the *FLO11* promoter when expressed as a cDNA clone in *Saccharomyces cerevisiae* (22). We selected *ffmA* for further study to examine its role as a regulator of *abcG1* expression.

### Phenotypes caused by loss of the *ffmA* gene

To confirm the effects of the *ffmA* gene on phenotypes, we prepared a new *ffmAΔ* allele in a different genetic background (AfS35). We also used the original disruption mutant library strain for comparison. Spores were prepared from the two wild-type strains and their isogenic *ffmAΔ* derivatives. A radial growth assay was performed for these 4 different strains (Figure 2A).

**Figure 2.**
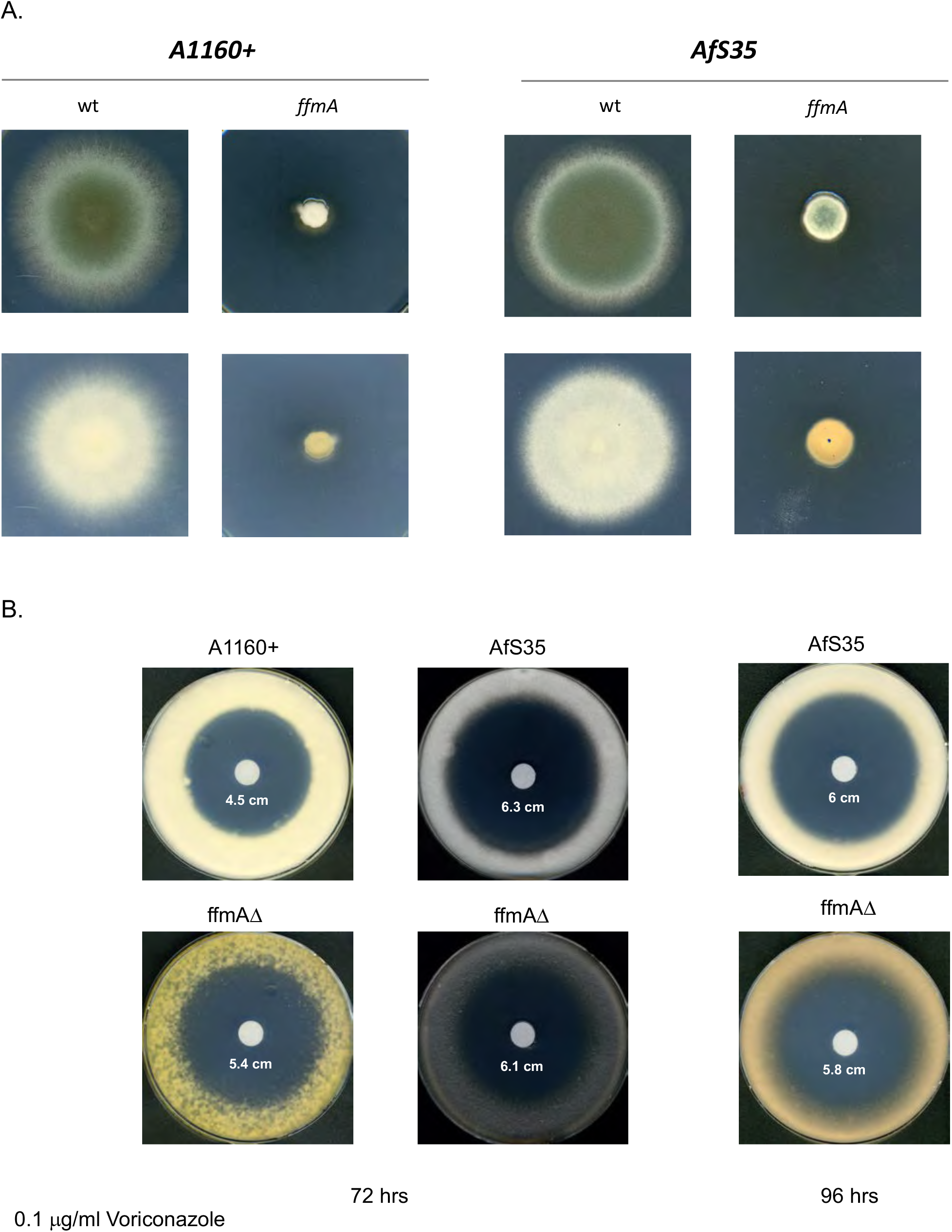
Phenotypes of *ffmAΔ* strains. A. The *ffmAΔ* strain present in the transcription factor disruption mutant collection (A1160 background) and a *ffmAΔ* strain constructed in the AfS35 genetic background were grown with their isogenic wild-type counterparts and spores isolated. Equal numbers of spores were placed on minimal medium and allowed to grow at 37°C. Note the orange color of the mycelial side of the *ffmAΔ* strains. B. Voriconazole susceptibility of a *ffmAΔ* strain. An equivalent number of spores from the strains indicated were spread on minimal medium and then a filter disk containing the indicated concentration of voriconazole was placed in the center of each plate. Plates were allowed to develop at 37°C for either 72 or 96 hours.

Irrespective of the genetic background employed, the clearest effect caused by loss of *ffmA* is a profound growth defect. We estimate the growth rate of the *ffmAΔ* strain to be roughly 25% that of the wild-type cells. This defect in growth rate complicated the analysis of azole susceptibility in the *ffmAΔ* strain. We also observed the accumulation of an orange/brown pigmentation upon loss of *ffmA* in both strain backgrounds. To compare the voriconazole susceptibility phenotypes of these strains, we used a zone of inhibition assay and placed a filter disk containing this azole drug in the center of plate of spores from each wild-type and isogenic *ffmAΔ* strain. These plates were incubated at 37°C and then photographed (Figure 2B).

Loss of *ffmA* from the A1160 wild-type strain caused a reproducible increased susceptibility to voriconazole but this was masked by the extreme growth defect seen in the AfS35 genetic background. The *ffmAΔ* derivative of AfS35 appeared to be more seriously growth compromised than its A1160 *ffmAΔ* counterpart.

Since loss of the *ffmA* gene caused such a pronounced defect in growth, we examined the effect of placing *ffmA* under control of a strong promoter. We hypothesized that overproduction of ffmA caused by replacement of the *ffmA* promoter with the corresponding region from the highly-expressed *hspA* gene might affect voriconazole resistance. Using CRISPR/cas9-mediated recombination, the *hspA* promoter was inserted in place of the native *ffmA* promoter region. Appropriate transformants were recovered and analyzed for several *ffmA*-dependent phenotypes.

In order to confirm that use of the *hspA* promoter to drive *ffmA* expression led to overproduction of FfmA, we assessed protein levels of ffmA using Western blot. To accomplish this, a rabbit polyclonal antiserum directed against bacterially-expressed FfmA was prepared. Isogenic wild-type and *hspA-ffmA* strains were grown with or without voriconazole treatment and levels of FfmA were analyzed by western blotting.

As shown on the left-hand side of Figure 3B, treatment of wild-type cells with voriconazole led to a 2-fold increase in FfmA levels compared to untreated cells. The presence of the *hspA-ffmA* allele caused a 2-fold increase in FfmA levels compared to the wild-type strain, consistent with a higher transcriptional level supported by the *hspA* promoter as we have seen before (23). Interestingly, exposure of *hspA-ffmA* cells to voriconazole caused a further increase in FfmA induction to roughly 5-fold above the azole untreated wild-type levels.

**Figure 3.**
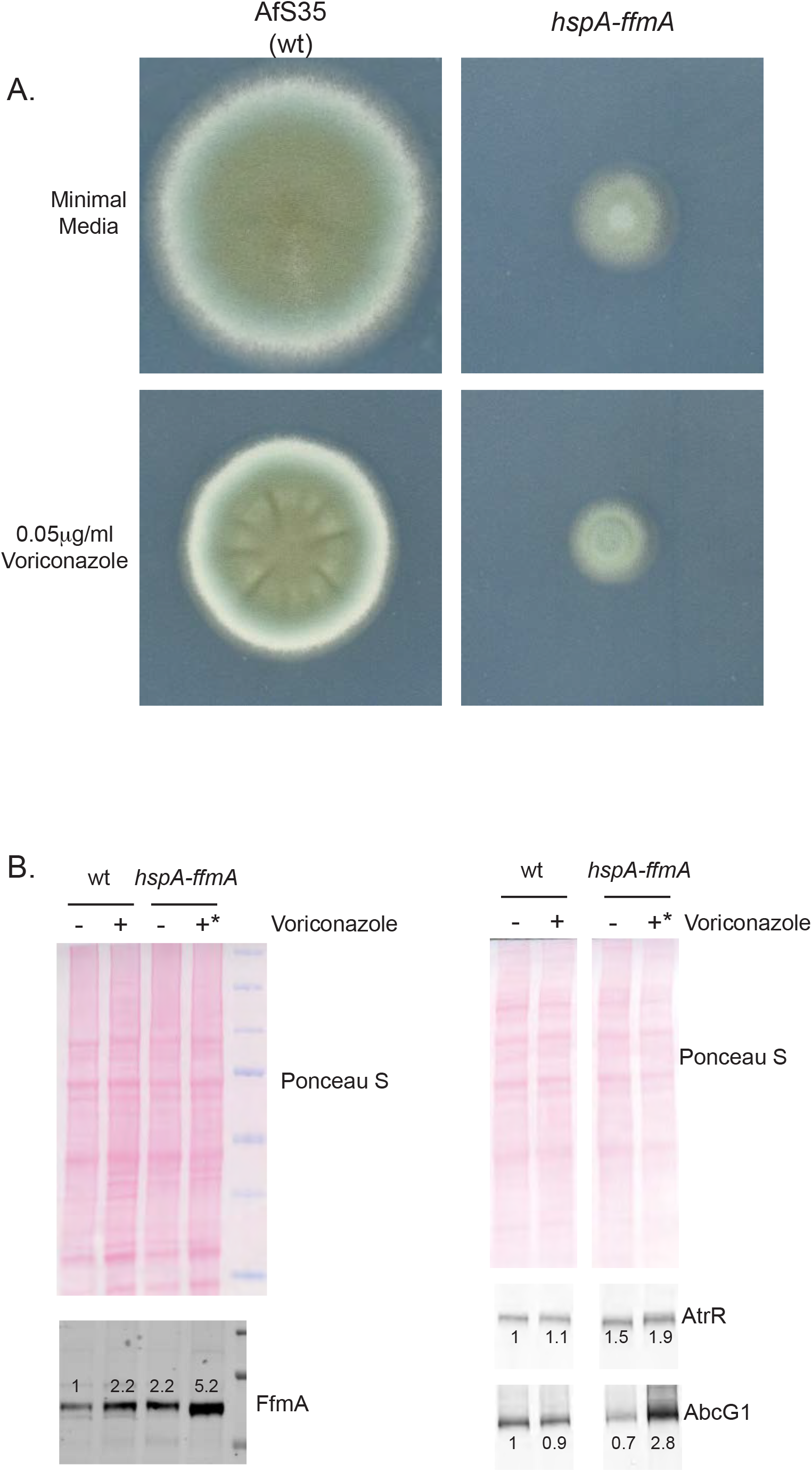
Driving *ffmA* using the strong *hspA* promoter causes a growth defect. A. The *hspA* promoter was inserted into the *ffmA* locus in place of the wild-type *ffmA* version. Equal numbers of spores from isogenic wild-type and *hspA-ffmA* strains were placed on minimal media without or containing 0.05 mg/l voriconazole. Plates were incubated at 37°C for 3 days and then photographed. B. Whole cell protein extracts were prepared from isogenic strains containing either the wild-type (wt) or *hspA*-driven (*hspA-ffmA*) *ffmA* gene. These strains were grown to mid-log phase in the presence or absence of 0.05 mg/l voriconazole. All strains were grown at 37°C for 16 hours except the *hspA-ffmA* strain in the presence of voriconazole that was grown for 24 hours due to its slow growth under these conditions. Equal amounts of extracts were analyzed by western blotting using the indicated rabbit antisera. The membranes were stained by Ponceau S dye to ensure equal loading and transfer. Western blot analysis using the anti-FfmA antiserum is shown in the left-hand panels while blotting with the anti-AtrR and anti-AbcG1 antisera are shown on the right. The numbers above each immunoreactive band refer to expression levels normalized to the wild-type in the absence of drug.

The presence of the *hspA-ffmA* fusion gene reduced radial growth to ~35% of normal 37°C (Figure 3A) similar to that seen for the *ffmAΔ* strain. This growth defect likely contributed to the lack of any effect seen on voriconazole susceptibility.

Our finding that loss of *ffmA* led to a decrease in AbcG1 expression (Figure 1) prompted us to examine the effect of *hspA-ffmA* on expression of both AbcG1 and the AtrR transcription factor that is key to the regulation of this ABC transporter-encoding gene (15, 16). The same protein extracts analyzed in Figure 3B were evaluated for the level of AbcG1 and AtrR in Figure 3C using appropriate antisera. Expression of AtrR was increased in the *hspA-ffmA* strain compared to wild-type by approximately 50% with a further increase to nearly 2-fold when the *hspA-ffmA* cells were challenged with voriconazole. Expression of the ABC transporter AbcG1 was seen to increase by 3-fold in *hspA-ffmA* cells in the presence of voriconazole compared to equivalently treated wild-type cells. These data further support the view that AbcG1 expression responds to changes in *ffmA* expression and suggest that the *atrR* gene might be another target of FfmA.

### Overproduction of AtrR can suppress ffmA-dependent phenotypes

Our finding of this potential link between AtrR and FfmA led us to examine the interaction between the two genes encoding these transcriptional regulators. To test the epistasis of *atrR* and *ffmA*, we used a hypermorphic form of *atrR* that we have characterized earlier: the *hspA-atrR* fusion gene. This allele of *atrR* was found to cause overproduction of *abcG1* along with other AtrR-regulated target genes (16)). We introduced the *ffmAΔ* null allele into a strain containing the *hspA-atrR* fusion at the normal *atrR* locus. Transformants were recovered and confirmed by PCR analyses.

The most obvious phenotype seen was the striking suppression of the growth defect of an *ffmAΔ* strain (Figure 4A). While the radial growth assay demonstrated that the *hspA-atrR* allele restored wild-type levels of growth to the *ffmAΔ* strains, production of orange pigment was retained indicating that this *ffmAΔ* phenotype was not suppressed. This indicates that while there is important overlap between FfmA- and AtrR-dependent phenotypes, this overlap is not universal between these factors.

**Figure 4.**
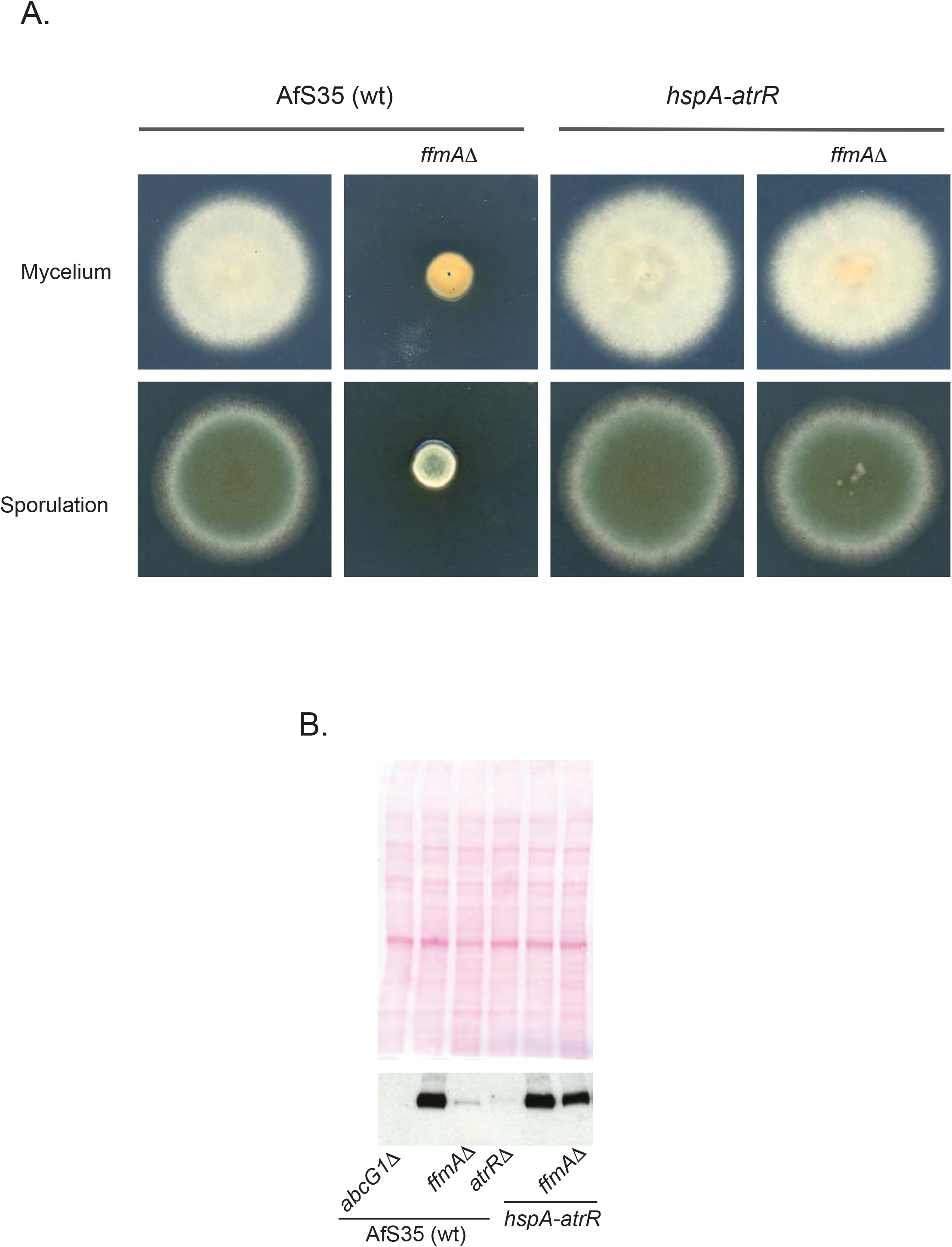
Epistasis of *atrR* and *ffmA*. A. To test the relationship between *atrR* and *ffmA*, the *ffmAΔ* cassette was introduced into a *hspA-atrR* fusion-containing strain. This double mutant strain was grown along with isogenic wild-type and *ffmAΔ* strains, spotted onto minimal medium and allowed to grow at 37°C. Plates were photographed after 3 days. B. Whole cell protein extracts were made from the strains described above and analyzed by western blotting using the anti-AbcG1 antiserum. The top panel shows Ponceau S staining of the membrane prior to blotting.

To examine the ability of the *hspA-atrR* fusion to suppress the defect in AbcG1 expression caused by the *ffmAΔ* lesion, we assessed levels of AbcG1 protein in these strains by western blotting (Figure 4B). Loss of *ffmA* decreased AbcG1 expression in an otherwise wild-type background while the *atrRD* null allele reduced AbcG1 expression even further and there was no detectable immunoreactivity in an *abcG1Δ* strain. Importantly, the *hspA-atrR* allele was able to restore AbcG1 expression to ~80% of wild-type in an *ffmAΔ* null strain. Overexpression of AtrR from the *hspA* promoter was sufficient to both restore near-normal growth and parental levels of AbcG1 expression to a strain lacking *ffmAΔ*.

### Doxycycline-dependent expression of *ffmA*

Both loss of *ffmA* and overproduction of FfmA from the *hspA* promoter caused a serious defect in normal growth along with reduction of both AbcG1 expression and voriconazole resistance. In order to develop a strain that could be grown and tested in a controllable manner we constructed a doxycycline-repressible form of *ffmA*. This was accomplished by integrating a doxycycline-repressible (dox-off; DO) promoter into the *ffmA* locus to replace the wild-type promoter region. This DO promoter was integrated either immediately upstream of the wild-type *ffmA* (DO-*ffmA*) or with a single Flag epitope fused to the N-terminus of FfmA (DO-Flag-*ffmA*). To confirm that the presence of the DO promoter did not cause a severe growth defect in the absence of doxycycline, appropriate transformants were placed on media containing or lacking this compound and allowed to grow at 37°C.

The presence of either DO allele of *ffmA* led to nearly normal growth in the absence of doxycycline (Figure 5A). The growth rate of the DO allele-containing strains was estimated to be 80% of the wild-type strain in the absence of doxycycline. The addition of 25 mg/l of doxycycline caused a reduced growth phenotype to 25% of the wild-type, resembling that of the *ffmAΔ* or *hspA-ffmA* strains analyzed above.

**Figure 5.**
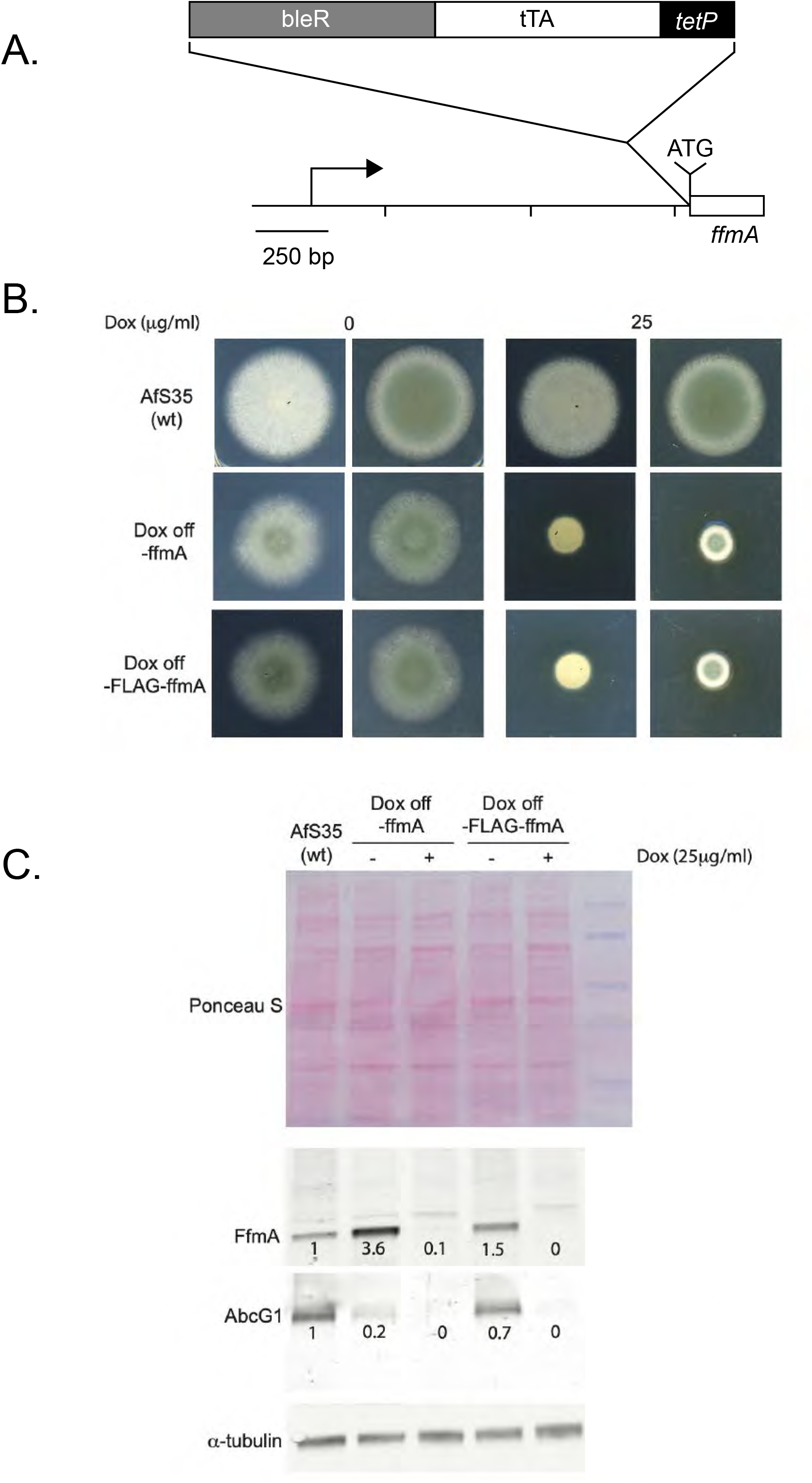
Conditional expression of FfmA using a doxycycline off promoter system. A. Schematic diagram for doxycycline off promoter integration into the ffmA gene. The structure of the doxycycline off promoter is shown by the bar at the top where bleR is the phleomycin resistance marker, tTA represents the region expressing the doxycycline off transactivator and tetP indicates the location of the doxycycline-responsive minimal promoter. When the FLAG tag is present, it is inserted between immediately before the native ffmA ATG (shown on the diagram). The *ffmA* gene contains a 1200 nucleotide long 5’ untranslated leader that starts at the position indicated by the rightward facing arrow. The scale of the diagram is indicated at the bottom left. B. The conditional doxycycline-repressible (dox off: DO) promoter was inserted just upstream of the ATG for the *ffmA*-encoded open reading frame, either with or without a single FLAG epitope fused to the FfmA amino-terminal methionine residue. This makes transcription of the DO-ffmA fusion gene sensitive to the presence of exogenous doxycycline. Isogenic wild-type and DO-ffmA strains were grown to mid log and then spores placed on minimal medium containing or lacking doxycycline at the indicated concentration. Plates were developed at 37°C as before. C. The strains above were grown in minimal media with no doxycycline to mid-log phase, then diluted into fresh medium lacking (−) or containing (+) the indicated concentration of doxycycline. Cultures were allowed to grow for 12 hours and whole cell protein extracts prepared. Equivalent amounts of each protein extract were analyzed for total loading, levels of FfmA and AbcG1 and tubulin using appropriate antibodies. The numbers below each specific immunoreactive band indicate expression level normalized to the wild-type strain.

We next used the doxycycline-repressible nature of the DO-*ffmA* and DO-Flag-*ffmA* genes to determine the effect of acutely repressing FfmA production on AbcG1. Both DO promoter-driven forms of *ffmA* were grown for 8 hours and then either left untreated or doxycycline added to the culture. After 12 additional hours of growth, whole cell protein extracts were prepared and analyzed by western blotting using anti-FfmA or anti-AbcG1 antisera (Figure 5B).

Both DO-driven forms of *ffmA* produced more FfmA protein than was observed in wild-type cells. FfmA levels were elevated to 3.6-fold higher than wild-type in the DO-*ffmA* fusion while the DO-Flag-*ffmA* fusion gene protein produced ~50% more FfmA protein than in wild-type cells. Surprisingly, AbcG1 expression was lowered in the presence of both DO-*ffmA* alleles when doxycycline was absent. The DO-*ffmA*-containing strain produced only 20% of normal AbcG1 protein while the DO-Flag-*ffmA* fusion led to production of 70% of wild-type AbcG1 levels. Both DO fusion genes were strongly repressed by the addition of doxycycline to the media with both FfmA and AbcG1 levels nearly undetectable. These data support the interpretation that loss of AbcG1 expression and the growth defect caused by loss of *ffmA* represent an authentic consequence of the absence of FfmA and are not an indirect effect caused by response to the growth defect elicited by FfmA loss.

### FfmA binds to the *ffmA* and *abcG1* promoters in vivo

The presence of a C_2_H_2_ DNA-binding domain in the FfmA protein sequence as well as its effect on *abcG1* and *atrR* expression is consistent with this protein playing a role as a transcription factor. To determine if FfmA might be directly associating with target gene promoters, we carried out single gene chromatin immunoprecipitation (ChIP) experiments using either the DO-Flag-*ffmA* fusion gene or wild-type cells. We used anti-Flag antibody to enable ChIP in the DO-Flag-FfmA strain and anti-FfmA in wild-type cells. Fixed chromatin samples from these two strains were sheared, immunoprecipitated with the appropriate antibody and then analyzed by qPCR.

ChIP using anti-Flag antibodies in samples from the DO-Flag-ffmA-containing strain showed strong enrichment of the Flag-FfmA protein on both the *abcG1* and *ffmA* promoters (Figure 6A). Note that in the case of this strain, the DO promoter was inserted between the normal *ffmA* promoter and the *ffmA* coding region, leading to retention of the wild-type *ffmA* promoter although it is no longer controlling expression of *ffmA*. We also examined ChIP of FfmA in unaltered AfS35 cells using anti-FfmA antibody. ChIP of FfmA in this strain showed similar enrichment of FfmA to both the fully-native *ffmA* and *abcG1* promoters. Both in the case of the Flag-FfmA and wild-type FfmA proteins, only marginal enrichment was seen at the *atrR* promoter, although it was greater than that seen on the actin promoter (*act1*) used as a presumptive negative control. These data are consistent with FfmA directly controlling gene expression of the *ffmA* and *abcG1* by direct promoter binding and potentially indirectly controlling *atrR* expression.

**Figure 6.**
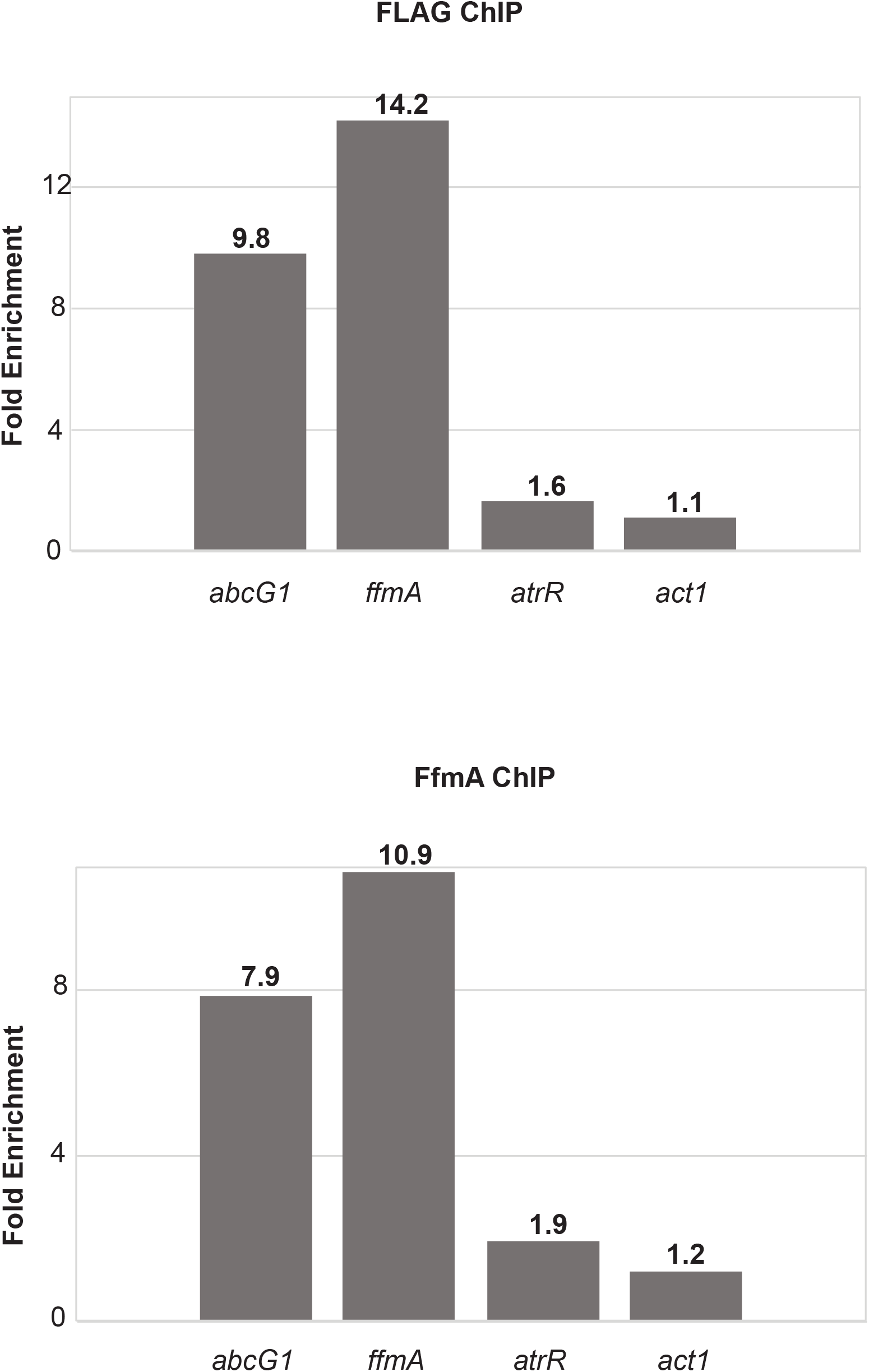
FfmA associates with promoter regions in vivo. Single gene chromatin immunoprecipitation experiments were carried out using either the DO-FLAG-*ffmA* strain (top panel) or wild-type AfS35 cells (Bottom panel). Strains were grown to mid-log phase in minimal medium and fixed chromatin prepared. After shearing, FfmA-bound DNA was recovered with anti-FLAG immunoprecipitation (Top panel) or anti-FfmA immunoprecipitation (Bottom panel). Immunoprecipitated DNA was analyzed using primer sets specific for the indicated 4 promoters. A control immunoprecipitation was performed with omission of the primary antibody as a control. The fold enrichment refers to the ratio of immunopurified DNA recovered in the presence of primary antibody compared to the absence. The numbers above each bar correspond to the fold enrichment from a representative trial. Note that *act1* immunopurification is used as a negative control promoter.

## Discussion

Our interest in *ffmA* emerged from the effect of this gene on itraconazole resistance as shown in the early analysis of a collection of transcription factor gene disruption mutants (17). More detailed analysis of the phenotype of a *ffmAΔ* strain clearly indicated that loss of this gene had a modest azole drug phenotype compared to its pronounced defect in growth. Other screening of this transcription factor disruption mutant library led to the detection of *ffmA* as a determinant required for resistance to cell wall stress agents but also showed a very similar general growth defect (19). Based on both our and the previous study of the phenotype of *ffmAΔ* strain, it seems that the primary phenotype caused by loss of *ffmA* is a severe growth deficit. This is in contrast to deletion of genes such as *srbA* (18) or *atrR* (15) that cause striking sensitivity to azole drugs but grow relatively well in drug-free media. FfmA is necessary for normal expression of *abcG1* but our data indicate a wider role in cell growth beyond azole resistance.

Based on the presence of a C_2_H_2_ zinc finger-containing DNA binding domain in its protein sequence, we believe that FfmA is a sequence-specific transcription factor. Our ChIP experiments support this view as FfmA is found to enrich on the *abcG1* promoter which would be consistent with FfmA acting to positively regulate expression of this gene. Additionally, we found that this same *abcG1* fragment was bound by AtrR (16), suggesting that FfmA and AtrR may bind the same or adjacent regions of the *abcG1* promoter We also found that FfmA showed enrichment on its own promoter, consistent with its expression being autoregulated.

The shared role for AtrR and FfmA in control of *abcG1* expression led us to examine the interaction between these two genes. Overproduction of AtrR using the *hspA* promoter was well tolerated by the cell and led to both increased azole resistance (16) and strong suppression of the growth phenotype caused by loss of *ffmA*. Owing to the difficulty in growing *ffmAΔ* strains, we disrupted *ffmA* in the *hspA-atrR* background. These transformants were readily obtained and grew normally. Furthermore, the contribution of FfmA to AbcG1 expression was strongly suppressed by the increased levels of AtrR. This genetic interaction led us first to suspect that FfmA might act upstream of AtrR in terms of function but ChIP experiments on the *atrR* promoter have not yet revealed strong enrichment for FfmA although we have not scanned the entire promoter. Another possibility is that both AtrR and FfmA stimulate expression of *abcG1* transcription and the increased levels of AtrR are adequate to reverse the transcriptional defect caused by loss of *ffmA*.

The precise role of FfmA in the cell remains to be determined. Clearly, *A. fumigatus* is exquisitely sensitive to modifications of FfmA expression. As we show here, either loss of *ffmA* or its overproduction from the *hspA* promoter caused a strong defect in normal growth. Driving either a Flag-tagged or untagged FfmA from the dox-off promoter did allow growth that was similar to wild-type but remained only 80% of normal. This sensitivity to variation in expression level has been noted before in the analysis of human disease genes (24) in which genes linked to disease are rarely seen to be associated with copy number variation owing to their critical importance. These genes are often associated with developmental processes as disturbances in the expression profiles of genes of this type lead to loss of viability. In the case of *ffmA* dosage changes, cells are still viable but clearly suffer significant impact on growth. Since this *ffmAΔ*-mediated growth defect can be robustly suppressed with *atrR* overproduction, we suggest that the downstream functions of these transcription factors may overlap. As discussed above, this is certainly true for control of *abcG1* expression but as *abcG1D* strains have no detectable growth defect in the absence of azole drugs, other target genes must be shared that impact growth under unstressed conditions. Identification of these genes will be an important future goal.

## Acknowledgements

We thank members of our labs for helpful discussions during the course of this work. This work was supported by NIH grant AI143198. We thank Dr. Jarrod Fortwendel for providing the dox off promoter and phleomycin marker constructs. Also thanks to Scot and Ryan Sargeant of Pacific Immunology for their excellence in production of polyclonal antisera.

## MATERIALS AND METHODS

### *A. fumigatus* strains, growth conditions, and transformation

The list of lab strains that were used in this study are listed in table 1. For initial shortlisting of strains that were defective for *abcG1* expression, the itraconazole-susceptible members of the transcription factor disruption mutant were used (17). *A. fumigatus* strains were typically grown at 37°C in rich medium (Sabouraud dextrose; 0.5% tryptone, 0.5% peptone, 2% dextrose [pH 5.6 ± 0.2]). Selection of transformants was performed using minimal medium (MM; 1% glucose, nitrate salts, trace elements, 2% agar [pH 6.5]). Trace elements, vitamins, and nitrate salts are as described in the appendix of (25), supplemented with 1% sorbitol and either 50 mg/liter phleomycin (after adjusting the media to pH 7) or 200 mg/liter hygromycin Gold (both InvivoGen). For solid medium, 1.5% agar was added. Doxycycline promoter shut off experiments were performed by adding 25 mg/liter doxycycline (BD Biosciences).

**TABLE 1.**
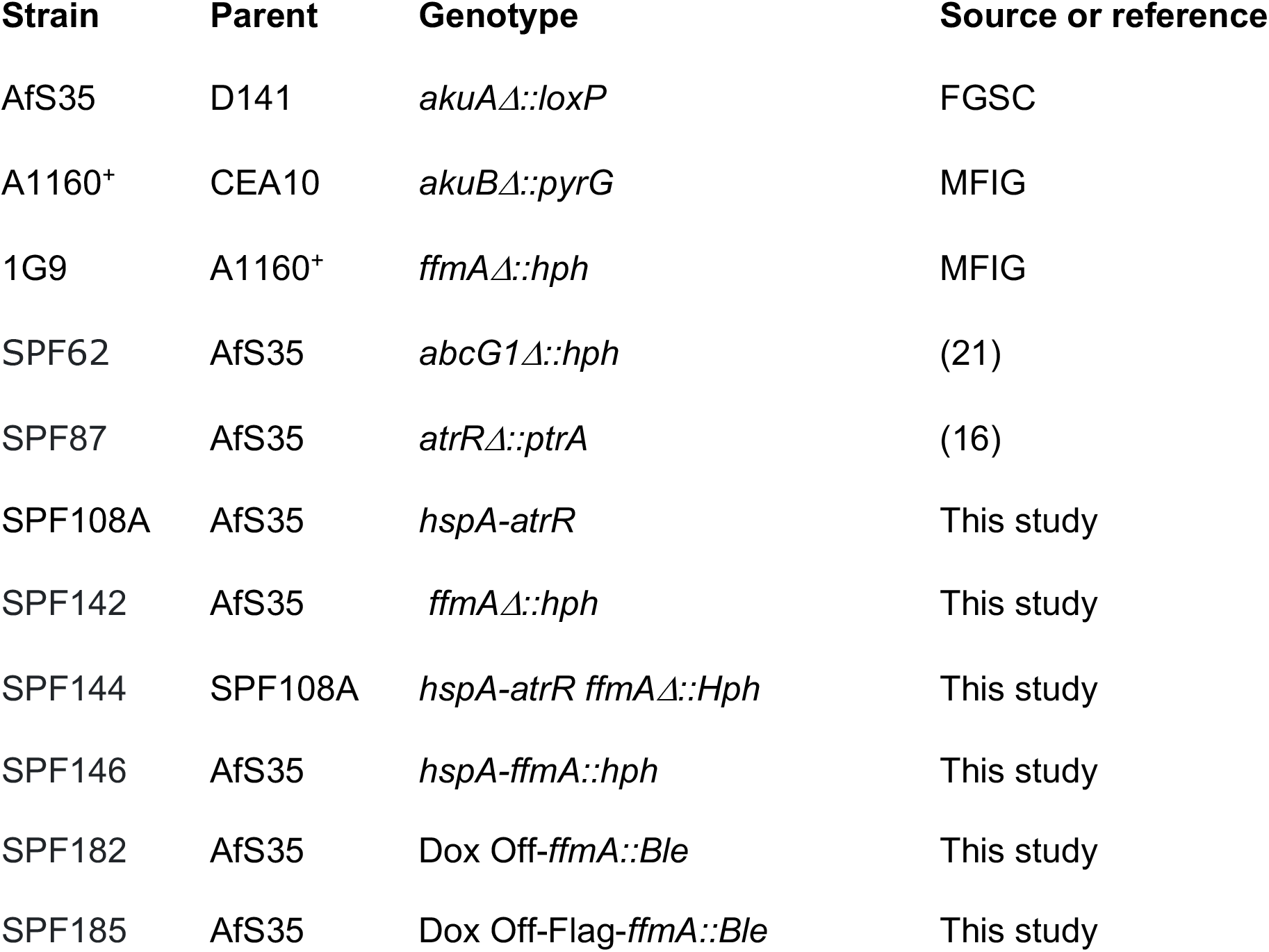
*A. fumigatus* strains used in this study

Transformation and generation of *ffmA* mutants was done using in vitro-assembled cas9-guide RNA ribonucleoproteins coupled with 50 bp microhomology repair templates (26). For generation of *ffmA* deletion mutants, 2 CRISPR RNAs (5’-ATCTAGGATCCATCATGAAG corresponding to the 5’ end of the gene and 5’-ATAGTCAATGGCTCAAGGGA corresponding to the 3’ end of the gene) were used to replace *ffmA* with the hygromycin resistance marker cassette amplified from the plasmid pSP62 (16) using ultramer grade oligonucleotides from IDT harboring 50 bp homology to the upstream and downstream junctions of *ffmA* gene. The doxycycline-repressible (Dox off:DO) promoter (with or without a Flag tag) marked with the phleomycin resistance cassette was inserted upstream of the *ffmA* gene using the single CRISPR RNA 5’-ATCTAGGATCCATCATGAAG. For generating the DO-*ffmA* allele, a microhomology repair template containing the resistance marker cassette and DO promoter were PCR amplified from plasmid pSP114. The plasmid pSP114 was generated from pSK606-*ptrA* (provided by J. Fortwendel), from which the *ptrA* marker was replaced by phleomycin resistant gene (Ble) by cloning the latter from pCH008-PhleoR (also from J. Fortwendel) at the KpnI & PstI sites. To generate the Ble-DO-Flag cassette, complementary oligonucleotides corresponding to a 1x Flag tag was annealed and ligated to the ends of PmeI-linearized pSP114. This plasmid was named pSP115, and was used as template to generate microhomology repair constructs to generate the Ble-DO-Flag-*ffmA* allele. Transformants were genotypically confirmed by diagnostic PCR of the novel upstream and downstream junction formed upon targeted integration, as well as by PCR amplification to confirm lack of the DNA binding domain of *ffmA* in the case of *ffmA* deletion mutants. At least 3 independent, properly-targeted transformants were phenotyped for all *ffmA* mutants, of which a representative strain is depicted in the data presented.

### Radial growth/drug disc diffusion assay

Fresh spores of *A. fumigatus* were suspended in 1x phosphate-buffered saline (PBS) supplemented with 0.01% Tween 20 (1x PBST). The spore suspension was counted using a hemocytometer to determine the spore concentration. Spores were then appropriately diluted in 1x PBST. For the drug diffusion assay, 1×10^6^ spores were mixed with 10 ml soft agar (0.7%) and poured over 15 ml of regular agar containing (1.5%) minimal medium. A paper disk was placed on the center of the plate, and 10 μl of either 1 mg/liter voriconazole was spotted onto the sterile filter paper. For the radial growth assay, ~100 spores (in 4 μl) were spotted on minimal medium with or without the drug. The plates were incubated at 37°C and inspected for growth every 12 h.

### Generation of an FfmA antibody

A region of 502 bp (corresponding to amino acids 195 – 340 based on homology to DNA binding domain of ScAdr1) from *ffmA* was PCR amplified and cloned in frame as a NdeI/HindIII fragment upstream of the C-terminal 6x-His tag in pET29a+ (EMD Millipore, Inc.) to form plasmid pSP113, and transformed into the *Escherichia coli* strain BL21(DE3). Two liters of transformed bacteria were grown to log phase and induced with isopropyl-β-D-thiogalactopyranoside for 90 min. Cell lysates were prepared using a French press and susequent protein purification accomplished using Talon metal affinity resin (TaKaRa Bio USA, Inc.) as described by the manufacturer. Protein fractions were analyzed by staining them with Coomassie blue and by Western blotting using His-specific antibodies. The purified proteins were then lyophilized and sent to Pacific Immunology (Ramona, CA) for injection into rabbits to generate polyclonal antibodies against FfmA. Antiserum generated from these rabbits was received and tested for immunoreactivity against *A. fumigatus* cell lysates. The antiserum was then purified using an AminoLink Plus coupling resin (Thermo Scientific, Inc.) according to the manufacturer’s instructions, and the affinity-purified antiserum was used to detect the FfmA protein from *A. fumigatus* cell lysates.

### Western Blotting

Western blotting was performed as described in (14). The FfmA polyclonal antibody used here has been detailed in the reference above, and was used at a 1:500 dilution. The anti-Flag M2 monoclonal antibody (F1804) was procured from Sigma, and used at a 1:2000 dilution. AbcG1 polyclonal antibody (21) was used at a dilution of 1:500.

### Real-time PCR

Reverse transcription quantitative PCR (RT-qPCR) was performed as described in (14), with the following modification: cell lysates were prepared from mycelial biofilm cultures formed upon inoculating 10^6^ spores in a petri dish containing 20 ml of Sabouraud dextrose broth and grown for 24 h at 37°C under nonshaking conditions. The threshold cycle (Ct) value of the *act1* transcript was used as a normalization control.

### Chromatin immunoprecipitation

Chromatin immunoprecipitation was done as described in (16) with the following modifications. 30 μl were reserved as an input control (IC) fraction for reverse crosslinking to verify sonication and control for ChIP and qPCR. The sheared chromatin was incubated with either anti-Flag M2 monoclonal antibody (Sigma-F1804) at 1:250 dilution or with anti-FfmA polyclonal antibody (referenced above) at a dilution of 1:50 overnight (16 h) on a nutator at 4^0^C. This sample was further incubated with 50 μl of washed dynabeads (Life Technologies) conjugated to either protein G (when using anti-Flag) or protein A (when using anti-FfmA) for another 8 h. Real-time PCR of ChIPed DNA was also performed as described in (15) with the following modifications. 0.5 μl of ChIP-ed or input (diluted 30-fold to bring it to 1%) DNA was used in 20 μl total volume reaction using SYBR green master mix (BioRad) and 0.4 μM of each primer. Fold enrichment was calculated to determine the enrichment of the promoter region for each gene. The oligonucleotide primer pairs used to check promoter enrichment consist of: cyp51A_F: 5’-GAGAAGGAAAGAAGCACTCTG, cyp51A-R: 5’-AgGAGGAAAATGGATAAGAGG; atrR-F: 5’-ACGGGATCCGTTTTGATACTC, atrR-R: 5’-CTGAACGAAGAGTCCGTCTC; abcG1-F: 5’-CGCTAATCATGAATCATCCCAC, abcG1-R: 5’-TCTCTTTTCTTGGACCCGAC; actA-F: 5’-GCCACCTAAGCGTTACCACT, actA-R: 5’-GCCGCTTCGTATAGGAGACC.

